# Social dilemma in the external immune system of the red flour beetle? It is a matter of time

**DOI:** 10.1101/128397

**Authors:** Chaitanya S. Gokhale, Arne Traulsen, Gerrit Joop

## Abstract

Sociobiology has revolutionized our understanding of interactions between organisms. Interactions may present a social dilemma where the interests of individual actors do not align with those of the group as a whole. Viewed through a sociobiological lens, nearly all interactions can be described in terms of their costs and benefits and a number of them then resemble a social dilemma. Numerous experimental systems, from bacteria to mammals, have been proposed as models for studying such dilemmas. Here we make use of the external immune system of the red flour beetle, *Tribolium castaneum*, to investigate how the experimental duration can affect whether the external secretion comprises a social dilemma or not. Some beetles (secretors) produce a costly quinone-rich external secretion that inhibits microbial growth in the surrounding environment, providing the secretors with direct personal benefits. However, since the antimicrobial secretion acts in the environment of the beetle it is potentially also advantageous to other beetles (non-secretors), who avoid the cost of producing the secretion. We test experimentally if the secretion qualifies as a public good. We find that in the short term, costly quinone secretion can be interpreted as a public good presenting a social dilemma where the presence of secretors increases the fitness of the group. In the long run, the benefit to the group of having more secretors vanishes and actually becomes detrimental to the group. Therefore, in such semi-natural environmental conditions, it turns out that qualifying a trait as social can be a matter of timing.

## Introduction

A social dilemma, such as the ‘tragedy of the commons’ (Gordon, 1954; Hardin, 1968), describes a situation where individuals have to prioritize between following a selfish action, which pays them individually in the short run, or an altruistic action such as maintaining a public resource, which will be beneficial for the group in the long run. In evolutionary biology, it is typically argued that natural selection will work against such altruistic actions unless a mechanism that promotes cooperation is in place (Hamilton, 1964; Nowak, 2006b).

Social dilemmas may occur under all kinds of social interactions. For example, whether or not one should get vaccinated (or have children vaccinated) against infectious diseases such as measles, mumps, and rubella is a case of a social dilemma (Bauch and Earn, 2004). Since vaccination might not prevent infection in 100% of cases and is not free of risk or side effects (Wu et al., 2011), the ideal situation is when everyone else would be vaccinated, as it prevents the spread of infectious diseases and allows an individual to avoid the risks and costs of individual vaccination. To understand the underlying dynamics of such dilemmas numerous mathematical models as well as experiments in various systems have been conducted (Sokoloff, 1972, 1974, 1977; Kerr et al., 2002; Nahum et al., 2011; Maclean and Gudelj, 2006; Nowak, 2006a; MacLean et al., 2010; Sigmund, 2010) some of which present resolutions to the dilemma either via ecological processes or by effectively changing the interactions.

Here we use *Tribolium castaneum* as a model system for inquiring whether the interactions resulting in a social dilemma are in fact consistent over time or merely as a result of when the observations were made (McCauley and Wade, 1981). The red flour beetle is used in various biological disciplines e.g. developmental biology, immunology, evolutionary ecology (Zou et al., 2007; Richards et al., 2008; Roth et al., 2010; Kittelmann et al., 2013; Joop et al., 2014) as a model invertebrate model organism. With short generation times of about one month, the beetles can be readily subjected to experimental evolution. Adult beetles have the innate immune defence, but in addition are capable of an external quinone-rich secretion with broad antimicrobial activity (Prendeville and Stevens, 2002). This secretion has been termed as an external immune defence (Joop et al., 2014) or alternatively, extended immune defence (Otti et al., 2014; Joop and Vilcinskas, 2014). Producing the external secretion is a genetically controlled trait and costly (Li et al., 2013; Joop et al., 2014) thus being a classic costly trait. This secretion can be toxic to the beetle larvae - and the secretors themselves - at high concentrations (Sokoloff, 1972; Joop et al., 2014). Flour beetles live in close proximity to each other and typically can be found in grain stores or flour mills, as human commensals, for thousands of years (Levinson and Levinson, 1994). Since the anti-microbial secretion acts in the environment of the beetle, it is potentially also advantageous to other individuals living in their proximity, including their offspring (Masri and Cremer, 2014). However, it remains to be tested if the secretion of the anti-microbial is actively controlled by the beetles and if it can be treated as public good, thus imposing a social dilemma in the beetles (Cotter and Kilner, 2010). If the beetles can indeed strategically modify their secretion of quinones to suit others and/or depending on environmental conditions, such as the presence of parasites or pathogens then this would provide conclusive evidence of the secretion being a social trait.

Due to above mentioned side effects of high concentrations of secretion, we hypothesise a population level upper threshold which if crossed, is potentially harmful to all offspring and the beetles themselves, resulting in reduced fitness. One would also expect a lower threshold, below which the sufficient spreading of quinones into the environment is no longer given to provide antimicrobial property. Hence, beetles would need to specifically control the amount they secrete into the environment. Since the secreted product can be utilised by other individuals in the population, secretion may be viewed as a social trait. The dilemma would be to either contribute to this common good and paying the costs or not to contribute, saving the costs and benefiting from the secretion of others. In the latter case, secreting beetles could compensate for non-secreting individuals in the population to secure an optimal quinone concentration in the environment.

In this study we address if quinone production meets the requirements of being a social trait. If populations of secretors are more successful at proliferating than populations of non-secretors and if in any mixed population, the non-secretors perform better than secretors, then quinone secretion could be classified as a social trait: secretion would increase the average fitness of the group at a direct cost to the providing individual. If there is no cost of producing and inhabiting a quinone rich environment then the average fitness of a group increases with the number of secretors and all secretor groups do better than a group of non-secretors. Here we tackle these two questions experimentally.

## Methods and Experimental Design

*T. castaneum* are convenient to breed in the lab, using a climate chamber (32 degrees C, 70% humidity), and population size as well as environment can be readily manipulated, e.g. by introducing parasites and pathogens. Overlapping generations are the natural scenario and therefore, all life stages are affected by the external immune defence being present in the environment. Hence, the costs of external secretion as well as its efficacy should manifest in the beetles’ fitness. Ideally, one would measure such as lifetime reproductive success. However, as the beetles can survive for several years under laboratory conditions, we took population growth as a proxy for fitness.

The environmental presence of the external secretion indeed protects the (non-secreting) beetle offspring from microsporidian parasitic infections within one generation of exposure (Joop et al., 2014). However, when exposed to the microsporidia over a longer time frame the beetles went extinct (Rafaluk et al., 2015). To avoid colony collapses here we used a fungus, *Beauveria bassiana* (B.b.), which upon single exposure is affected by the external secretion of the beetles while in the long run becomes resistant and more virulent (Rafaluk et al., 2017). Here we chose a fungus spore concentration of 10^8^ spores per gram of flour for the preparation of the spore-flour mixture (Rafaluk et al., 2017). As a stress control and to control for host-parasite interactions, we also introduced a treatment using heat killed spores of the fungus. Therefore, we had three levels of pathogen (none/control, non-infecting (heat killed) *B. bassiana* and infecting *B. bassiana*). We investigated whether

- on an individual level, secretors compensate for non-secretors
- the average fitness of the group increases with the number of secretors, and
- the fitness of populations with only secretors is greater than the fitness of the popula-tions with no secretors.

To address these topics we first needed to define secretors vs non-secretors at an individual level since the beetles produce different levels of quinones. Within the “non-secretors”, individuals produce very low levels of quinones, but there are none that do not produce any. For each treatment, we had one population, which was composed of all low level secretors. The level of quinone secretion which this low level secretor population reached in each particular treatment was taken to be the baseline quinone secretion for that treatment (Fig. 1). Quinones of interest auto-fluoresce with a peak at 245 and 246 nm. To fully capture the quinone secretion on a plate reader we took measures from 210 to 262 nm and calculated the area under curve (Joop et al., 2014; Rafaluk et al., 2017). As a conservative estimate all the individuals secreting more quinone than twice the baseline secretion were considered to be secretors, for that treatment.

**Figure 1:**
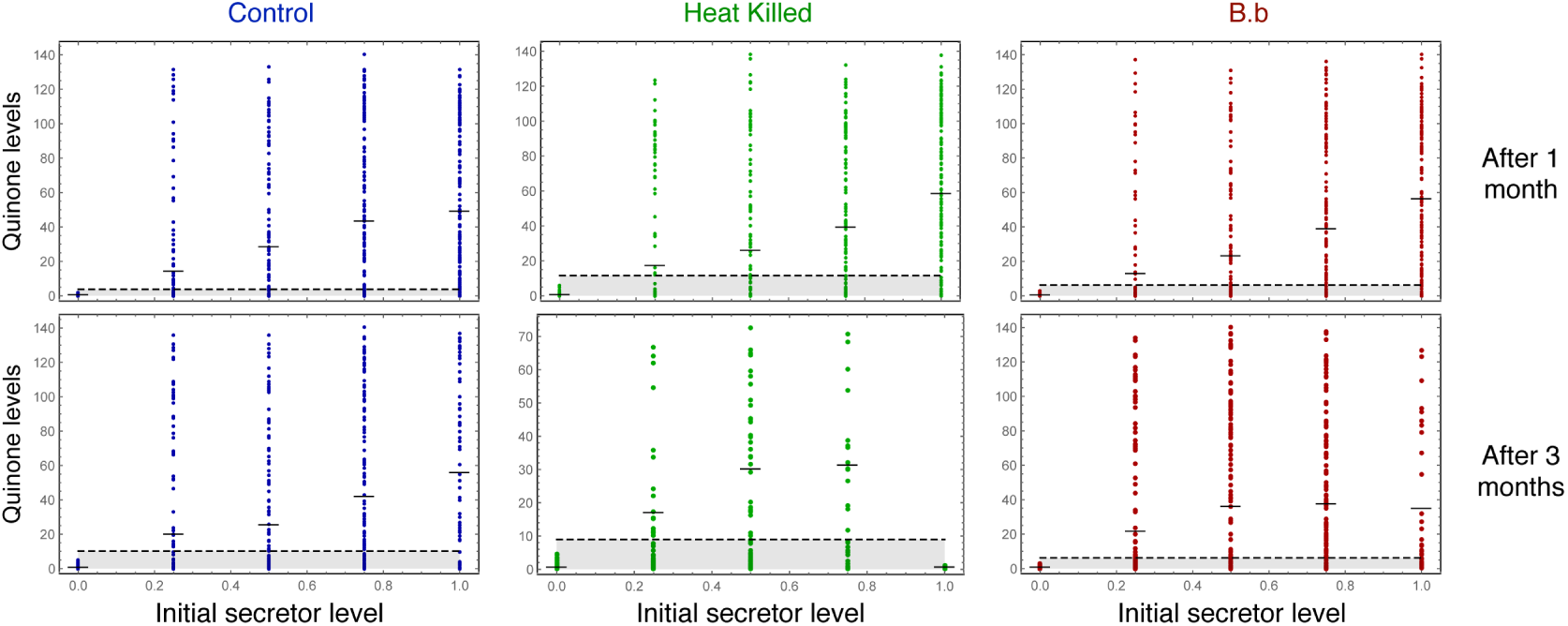
Defining secretors vs. non-secretors. The individual quinone levels where assessed for 30 randomly sampled individuals from each replicate are represented as a dot. In total the aim was to get 150 individuals from each treatment. In some cases, the population size was not large enough for this measurements. Even in cases where there are no secretors, there is a basal level of expression. As a conservative assumption we set a level twice that of the maximum basal expression (given by the horizontal dashed line) to differentiate between secretors and non-secretors. We then extrapolate this measure to the population sizes given in Fig. 3.

To address the point of compensation for the presence of non-secretors we first needed to establish a baseline so as to see what a non-compensatory scenario looks like. From the parental population which was used to seed the different replicates, the mean level of quinone secretion of the secretors was *q*_*s*_=57.74 (*n*=46) (measured as above) and that of the non-secretors was *q*_*ns*_=1.82 (*n*=46). We normalised these means such that they formed the end points of a continuum from 0 to 1 ((1.82−*q*_*ns*_)*/*(*q*_*s*_− *q*_*ns*_) to (57.74−*q*_*ns*_)*/*(*q*_*s*_− *q*_*ns*_)). Thus a group with only secretors coming from this population should theoretically produce quinones at level 1 and conversely all non-secretor groups at level 0. Additionally assuming a linear accumulation of quinones, a group made up of *i* secretors and *N*−*i* non-secretors should produce *i/N* level of quinones (again scaled between 0 and 1).

We manipulated the initial ratio of secretors to non-secretors in a population with a given population density and carrying capacity. Here, we used one beetle per 0.1g flour, 60 beetles per population in 6g flour (Blaser and Schmid-Hempel, 2005). The beetle lines originated from an evolution experiment on quinone secretion, selecting for either high (= secretors) or low (= non-secretors) quinone secretion (Joop et al., 2014). We implementd five ratios of secretors:non-secretors (100% secretors, 75:25, 50:50, 25:75, 100% non-secretors), each level being replicated five times. Thus the experiment was performed as a fully factorial set up, resulting in a total of 75 populations per time point, with the time points being one month and three months.

As the beetles influence their environment due to their external immune secretion as well as other excretions, the environment is not constant in our experiment but changes over time. The secreted quinones spread, oxidise to hydroquinones which also have antimicrobial property, and persist. Therefore, populations started in quinone free environment (with or without pathogens, depending on the treatment), while later generations as well as ageing adults experienced a quinone rich environment, enriched with additional excretions and most likely containing less food or food of lower quality (compare (Joop et al., 2014) for quinone flux into the environment). As mentioned, *T. castaneum* can live under laboratory conditions for several years. In the wild however, lifetime is expected to be shorter. To consider the changing environment we measured the population growth at two time-points, after one month and after three months, representing a likely natural life span of an adult and allowing for about three generations in the three month treatment including the starting generation. The groups that were set up had been randomly assigned to a one month or three month treatment. This approach avoided the problem of disturbing the *Tribolium* jars after a month (including the danger of eggs/larval wounding during sieving, changing the flour structure etc.).

At the end of the assay, i.e. after one or three month all individuals per jar were counted and separated by life stage. In addition, quinone secretion was measured for thirty randomly selected individuals to calculate the mean quinone level (compare (Joop et al., 2014)). In cases where less than thirty adults were present in the population, all remaining adults were measured.

## Results

### Secretors compensating for non-secretors

We indirectly tested if secretors compensate for non-secretors by comparing the quinone level after the experiment to the theoretical quinone level expected if the beetles did not compensate for non-secretors. For example the 75:25 secretors:non-secretors populations should have about 75% of the quinone secretion of an all secretor population within the given treatment. We could come to the conclusion that secretors compensate for non-secretors if the three different secretors:non-secretors populations would show significantly higher quinone secretion than expected from this calculation. To assess the increase/decrease in the level of quinones secreted by the beetles first we established a baseline which could be compared to the theoretically expected value as defined in the Methods section. The actual amount of quinone secreted in each cohort of different fractions of secretors was measured and again rescaled as per the Methods to fall between 0 and 1 defined by the parental generation. The data points in Fig. 2 are not at the initial fraction of secretors as we take into account the correction due to the changed fraction of secretors over time. We find only minimal deviation from the expected amount of quinone (Fig. 2). The secretors did not over (or under) compensate for the presence (absence) of non-secretors. Thus the individual quinone production does not seem to be under any active control by the beetle itself.

**Figure 2:**
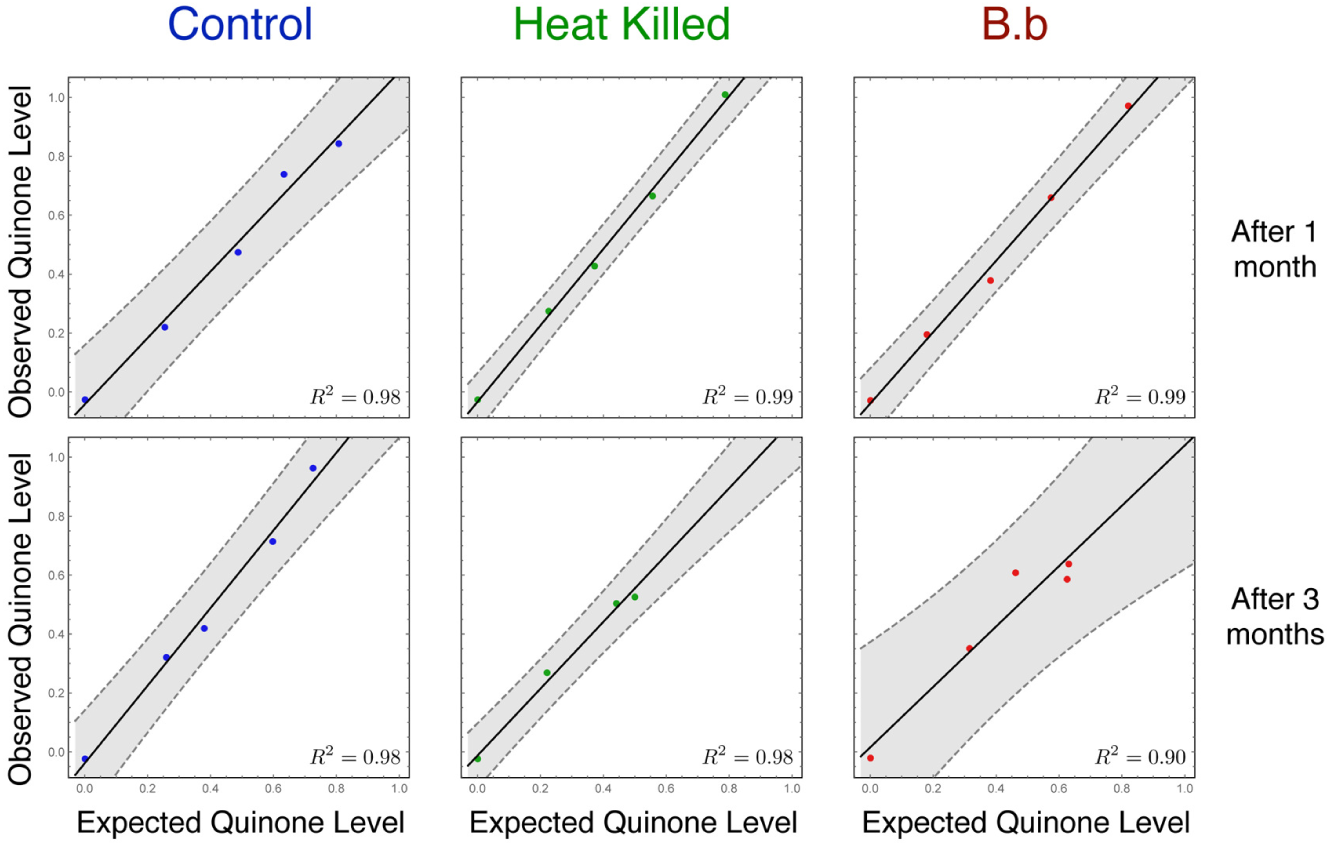
Comparing the observed quinone levels from the experiment with the expected levels calculated under linearity. If the secretor do not compensate for the non-secreting beetles, then the amount of quinone expected in the populations would be linear as to the fraction of secretors in each population. The quinone levels lie between 0 and 1 as forming the ends of a continuum where 0 corresponds to *q*_*ns*_=1.82, the mean quinone level of a non-secreting population and 1 to *q*_*s*_=57.75, the mean quinone level of a secreting population as from the seeding parental generation (see Sec. Results). Note that the expected quinone levels (fraction of secretors) are not constant as (0, 0.25, 0.5, 0.75, 1) but change with the fraction of secretors after one and three months as a result of the population dynamics (Fig. 4). The expectation is based on the assumption that the production of quinones is unaffected or cannot be actively modulated by the secretor in response to the presence/absence of non-secretors The strong linear relationship between the shows that the secretors do not compensate/modulate their secretion for the presence of non-producers. The solid lines are a linear fit to the data with a 95% confidence region shaded around it.

### Population growth

We take the growth rates of the populations as proxies for fitness. If the secretion of quinones is a public good then the fitness of the group would increase with the number of secretors in it. Thus the all secretor groups would have the highest fitness - population size - of all. After one month, the fitness of the populations increased with the fraction of secretors increases, in all three treatments (Fig. 3, One-month). Within most of the mixed populations the fraction of secretors decreases over time i.e. they generally do worse than non-secretors (Fig. 4). Thus, this points to the existence of a social dilemma based on the expression of quinones. Note that in contrast to the general trend, the population trajectories starting at 0.5 in B.b and heat-killed treatments and at 0.25 in all three treatments, show either minimal deviation from the starting fraction (Fig. 4).

**Figure 3:**
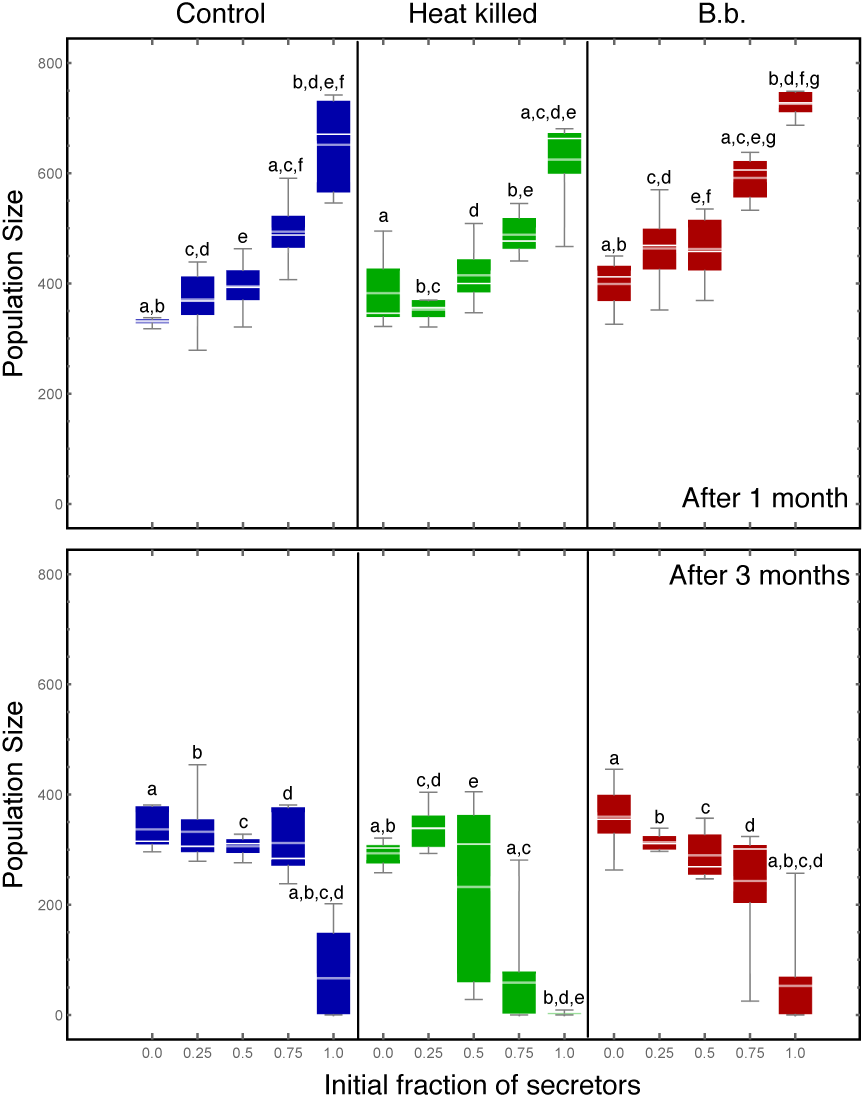
Population size dynamics in the three treatments for the different initial ratios of secretors:non-secretors. In the first month, the all-secretors group achieves the largest population size in all three treatments, with the growth being monotonic in the number of secretors at the initial conditions. After three months however in all three treatments, the all-secretors suffer population crashes. This could be due to the population reaching its ecological limit the earliest as compared to others due to the initially fast growth. Also, it is possible that the amount of quinone produced cannot be controlled individually by the beetles and thus a population crash results due to excessive quinones becoming toxic. Within each treatment (and control) panel we performed an ANOVA between the distributions. The final increase (after one month) or decrease (after three months) is always significant. For the control, heat killed and B.b. treatments after 1 month, the results were significant at *p* < 0:05 level F_4,20_ = 22:39, *p* = 3:7 × 10^−7^, F_4,20_ = 22:39; *p* = 4:8 × 10^−6^ and F_4,20_ = 22:39, *p* = 5 × 10^−8^ respectively indicated by like letters. After three months the decrease in population size is significant again with F_4,20_ = 16:71, *p* = 3:6 × 10^−6^, F_4,20_ = 11:49, *p* = 5:2 × 10^−5^ and F_4,20_ = 10:04, *p* = 1:2 × 10^−4^ for control, heat killed and B.b. treatments respectively.

**Figure 4:**
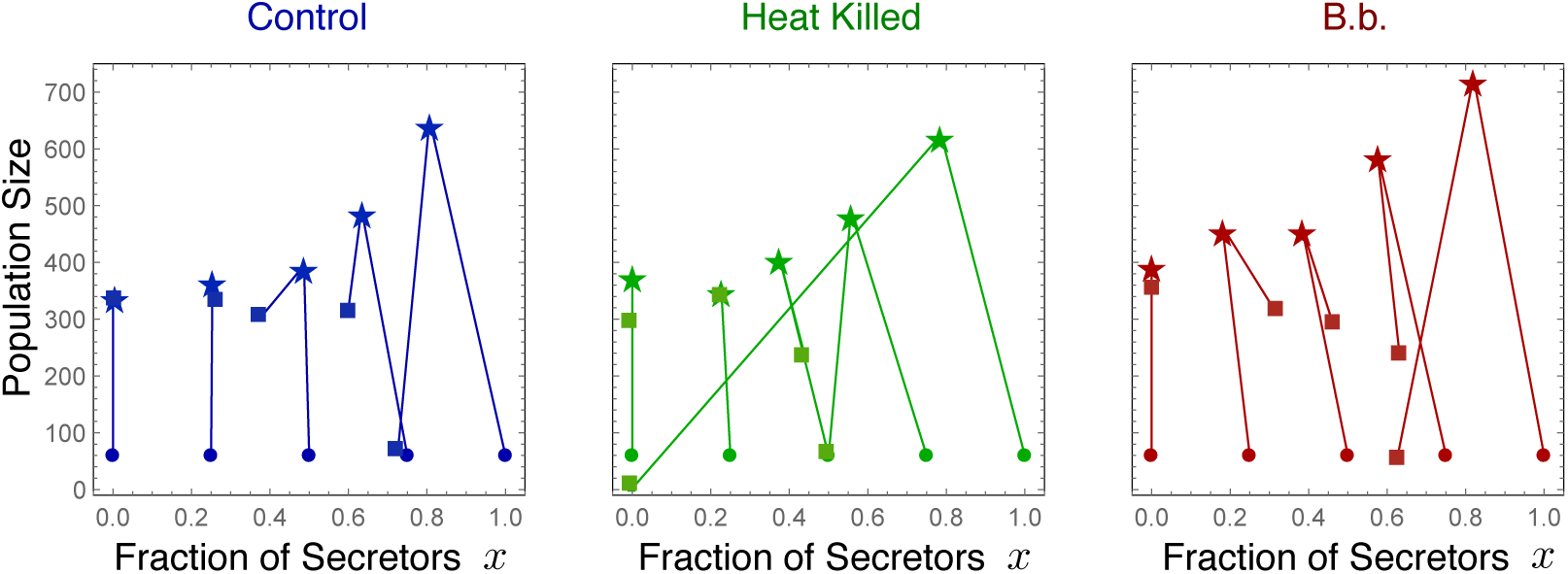
Population dynamics is usually excluded from the analysis of evolutionary dynamics. The experiment begins at the points indicated by circles, the one month time point represented by stars and the three month time point is given by squares. However when we look at ecological and evolutionary dynamics together we see that all treatments experience a growth in population size, which reaches around 300 individuals in almost all populations except with all secretors (and the one with 75% secretors in the heat killed treatment) before reducing. This might indicate that for the given environment a population limit is reached with about 300 individuals. Furthermore, the more initial secretors the stronger the drop down, potentially indicating that the upper quinone-threshold was crossed. This in combination with expending the environment finally results in extinction, whereas populations which start with less secretors grow slower and thus still survive after three months.

If we instead focus on the results after three months, the interpretation is very different (Fig. 3, Three-months and Fig. 4). After three months, the fitness of the population decreases with the initial fraction of secretors, implying no social dilemma between secretors and non-secretors.

### Population dynamics

An important difference between different time points is the population dynamics: population size increased rapidly over the first month, but after three months the population had declined substantially again (Figs 3, 4). It seems that effects of the selection procedure and stochasticity become more pronounced only later (Fig. 4). All populations decreased in size after three months. However, the fraction of secretors at both time points, in most of the treatments, decreased, indicating that non-secretors are advantageous. This again hints at a social dilemma, but the slowed-down growth of populations with more secretors after three months clearly shows that one cannot speak of a social dilemma in this system in general.

As for the population sizes, specifically, after three months with the different initial conditions we found:

• 0% secretors - By definition there are no secretors in all three treatments for this initial condition. For B.b. and the control treatment the final populations were around 350 while for the heat killed version they were slightly lower around 300.

• 25% secretors - The final population size in all three treatments stabilised at around 300. The fraction of secretors stabilised to around 20-25% for the control and the heat killed treatments but with active pathogen pressure in the B.b. treatment it increased to about 35%.

• 50% secretors - For the B.b and the control, the final population sizes were close to 300. The final population size was slightly lower for the heat killed treatment. The final secretor levels for the two pathogen treatments stayed almost close to 50% while for the control it dropped to 40%.

• 75% secretors - We observed a major population crash in the heat killed treatment with the fraction of secretors close to 50%. This was followed by the B.b. treatment which reached a population size of 250 with 65% of secretors while the control had a higher population size of 300 also with about 60% of secretors.

• 100% secretors - The heat killed treatment again suffered from the most drastic population crash with the fraction of secretors along-with. For the B.b. treatment the population also decreased in size but the fraction of secretors remained substantial at 60% while in the control the fraction of secretors was close to 70% albeit in a small population.

While in the short term (one month) all-producer populations did best, in the long term (three months) mixed populations with less than 50% secretors appeared to be more stable (Fig. 3, 4). This could arise from the limited environment, where quinone levels aggregate beyond the optimal level. Most importantly, this again proposes a fixed rather than a flexible strategy in quinone secretion. Seemingly, individuals are not adapting their secretion to environmental parameters such as pathogens or population composition or environmental quinone concentration, but the ratio of secretors to non-secretors may be underlying the maximal group benefit. This is also reflected in the change in the population sizes, where in the initially all-producer cases growth rate drops to almost zero after an initial boost (Fig. 3).

### Discussion

Our study shows that it is not possible to conclude that an observed “social” trait presents a social dilemma in general. It might very well be possible that the trait that seems social, might be a simple by-product of a natural phenomena one which cannot be controlled. In case of the external immune system of *Tribolium*, are other beetles getting a benefit from the producer beetles just as a by-product? In that case it would not be a social trait. This depends on the cost of producing the antimicrobial products. While it has been previously shown that indeed it is costly to produce quinones (Li et al., 2013; Joop et al., 2014), this does not mean that it would be necessarily amenable to manipulation by the secreting beetles themselves. Herein we show that population density, environmental concentration, pathogen presence and the proportion of secretors vs. non-secretors do not trigger an active regulation of quinone production in *Tribolium castaneum*.

We show that in a very simple setup a social dilemma can be found after one month but if we look at the all the results together and not just only population dynamics or only trait dynamics then after three months there is no sign of a social dilemma. Thus the time point at which experiments are studied can be crucial in determining whether a social dilemma is present or not. Extrapolating an observation as a life history trait, can resulting to misleading conclusions about experimental and empirical systems. Recently, for example *Pseudomonas* bacteria came under scrutiny for similar reasons. Interpreting any external secretion as a cooperative trait is misleading where every interaction could then be cast in the sociobiological framework (Foster et al., 2007; Nadell et al., 2009; West et al., 2006; Rainey et al., 2014). An external secretion can be interpreted as a public good in a certain case while not in others and this can depend on the genotype-environment interaction (Zhang and Rainey, 2013; Rainey et al., 2014). Thus, where the natural environment of an organism is not provided, any desired result could potentially be obtained by manipulating the environment. Yeast is another example where the environment can easily produce an interaction pattern capable of being captured by a social dilemma of one form or the other: Invertase produced by some yeast cells hydrolyses sucrose into glucose and fructose which are then available for uptake. However, in natural populations, not all yeast cells produce invertase. Thus avoiding the invertase production costs, the non-secretors can uptake the simpler sugars owing to the presence of the secretor yeast cells. While it is tempting to classify this system as an example of a social dilemma, over time as new data has emerged, the perceptions have changed (Greig and Travisano, 2004; Gore et al., 2009; MacLean et al., 2010). What was thought of as a tragedy of the commons was then interpreted as a snowdrift game just to be later demonstrated as not to be a social dilemma at all.

In a similar fashion here we show that it is possible to interpret the *Tribolium* system as appropriate to study social evolution (in particular, social dilemmas) or not, depending on the duration of the study. The potential common good in this system would be the quinone secretion with its antimicrobial property, being produced by adults only but with potential benefit to all life stages. Furthermore, *Tribolium* beetles are capable of mate choice (Peuß et al., 2015) as well as of kin-recognition (Jasienski et al., 1988; Parsons et al., 2013). Should it be the case indeed that there are no metabolic costs, rather than we have not found them so far (thus resulting in cheating), an alternative explanation may be provided by actively regulating the population composition, e.g. by cannibalism or differential dispersal rates between secretors and non-secretors. Under environmental conditions requiring less external protection, such as a parasite-free environment, there would be no need to take on the toxicity of quinone secretion and non-producers might cannibalise producer-offspring, while in a contaminated environment they would not do so to ensure the sufficient quinone supply for their own offspring. So ‘cheating’ would be shifted to a different level, which remains to be shown.

However, *Tribolium* species are not the only insects to secrete quinones. Bombardier beetles use it as predation defence, spraying it towards the predator to fend it off (Eisner et al., 1977), an interesting trait, in a solitary beetle. On the other hand, earwigs, sub-social insects providing facultative maternal care (Wong and K¨olliker, 2012) and showing sibling cooperation (Falk et al., 2014), can produce and secret quinones in the larval as well as the adult life stage (Gasch et al., 2013; Gasch and Vilcinskas, 2014). For this species quinone secretion has been discussed as predation defence in larvae (Gasch and Vilcinskas, 2014), while adults also use it when applying parental care to their eggs (Meunier and K¨olliker, 2012). These examples point towards the actual ability of quinone production being rather evolutionary conserved, while its utilisation seems to be diverse. Future research therefore not only should address the evolution of sociality as such, but considering it to be a gradient sensu lato (Cotter and Kilner, 2010).

Furthermore, our results reflect on how well experimental conditions represent the natural scenario. In case of pathogen free conditions, it is possible that the secretion decreases to extremely low levels or might even be lost. We rely on the pathogen condition to test our hypothesis and do not consider long term dynamics, which should help to alleviate this potential issue (Mitschke et al. unpublished work) (Joop et al., 2014). Competition for food is another likely scenario persisting under various conditions, staying in a toxic environment appears to be less likely at least when considering motile organisms. For these, the solution to both problems would be dispersal, which is not permitted in most experimental setups. Thus, systems with time-dependent conditions put us at risk of generating results fitting a theoretical expectation rather than objectively studying a natural system in an experiment. This stresses the importance of designing experiments which are informed but not constrained by the current theoretical thought before making final conclusions about the social interactions in the system.

## Acknowledgements

Thanks to T.G. Mohr and C. Rafaluk for assisting in the lab. This project was funded by a Volkswagen advanced postdoctoral grant to GJ (87133). CSG and AT thank the generous funding from the Max Planck Society. The authors thank David W. Rogers for improving the clarity of the manuscript. The authors declare no conflict of interest.

